# Integrated Isoform-Resolved Transcriptomic Analysis of *Gossypium barbadense* and *Gossypium hirsutum* Fibers

**DOI:** 10.1101/2024.10.11.617711

**Authors:** Jiwei Tang, Xinxin Gu, Yue Ma, Qingying Meng, Peihao Xie, Shihe Jiang, Liuyang Hui, Yiyang Lang, Mengqin Tang, Ying Zheng, Daojun Yuan

## Abstract

A comprehensive understanding of RNA expression and splicing during cotton fiber development plays a crucial role in explaining the differences in fiber quality between two different cotton species. To date, most cotton transcriptomic studies have utilized short-read sequencing data, which limits the ability to capture full-length mRNAs. In this study, we compiled long-read RNA sequencing data from the PacBio platform, as well as short-read RNA sequencing data from 10 fiber developmental stages, for both *Gossypium barbadense* and *Gossypium hirsutum*. We identified 183,767 and 178,994 isoforms in *Gossypium barbadense* and *Gossypium hirsutum*, respectively, generating the most comprehensive transcriptomic atlas of cotton to date. Alternative splicing events exhibited stage-specific variations during fiber development, and co-expression network analysis, combined with LASSO regression, identified isoforms highly correlated with each developmental stage. These findings reveal previously uncharacterized aspects of isoform regulation during fiber development and provide valuable resources for further research into the molecular mechanisms underlying fiber quality differences between cotton species.

## Background

Cotton is a vital and multifaceted economic crop with applications extending well beyond textiles, apparel, and healthcare industries. What is less known, however, is its use in the production of paper currency. Specifically, cotton fibers constitute the principal material in the manufacture of RMB.

Cotton fibers originate from the epidermal cells on the ovule, known as trichomes (Wu et al 2018), The development of these fibers is a complex process that is divided into five distinct stages: (1) Initiation and Differentiation (-3 to 5 DPA), where ovule epidermal cells proliferate and start to differentiate into fiber cells under the regulation of gibberellins and auxins; (2) Primary Cell Wall Synthesis (5-16 DPA), characterized by the production of a cellulose-rich primary cell wall; (3) Transition from Primary to Secondary Cell Wall (16-20 DPA), marked by a shift towards synthesizing secondary cell walls that contain increased amounts of lignin and hemicellulose; (4) Secondary Cell Wall Synthesis (20-40 DPA), a key phase for determining fiber quality through the development of the thickness and composition of the secondary cell wall; (5) Maturity (40-50 DPA), during which fiber cells cease growth and the synthesis of the secondary cell wall gradually ends(Tian and Zhang 2021).

The primary components of secondary cell walls are cellulose, hemicellulose, and lignin. During the development of these cell walls, GhKNL1, a transcriptional repressor, plays a crucial role in the development of cotton fibers. It binds to GhCesA4-2/4-4/8-2 and GhMYB46 to regulate the synthesis of cellulose. Additionally, GhKNL1 interacts with the promoters of GhEXPA2D/4A-1/4D-1/13A, inhibiting the expression of these genes and thus modulating fiber elongation(Wang et al 2022).

In eukaryotes, the modulation of gene expression involves the generation of various isoforms through differential splicing, which subsequently translate into distinct proteins, manifesting diverse phenotypes that facilitate adaptation to environmental changes. While the majority of transcriptomic analyses focus on gene-level expression, assessing the regulatory impact of gene expression throughout developmental processes, it is crucial to recognize that individual genes can produce multiple isoforms via alternative splicing. These isoforms often exhibit significant differences in structure and function, and isoform-level investigations can unveil phenomena that remain undetected at the gene level. It is pertinent to clarify that in numerous studies, the terms ’isoform’ and ’transcript’ are frequently used interchangeably. Strictly speaking, an isoform represents one of the multiple transcripts that are produced by the same gene through distinct splicing events. Conversely, a transcript specifically refers to the mRNA that is actually expressed. Thus, the term ’isoform’ reflects the intrinsic diversity within a single gene, whereas ’transcript’ denotes the specific RNA products generated during the gene expression process.

Alternative splicing is a post-transcriptional regulatory mechanism in which DNA initially transcribes pre-mRNA that undergoes RNA splicing, involving the excision of introns and the ligation of exons, to form mature mRNA (also referred to as a transcript isoform), which includes 5’ and 3’ untranslated regions (5’/3’ UTRs) and a coding region. The phenomenon of alternative splicing was first discovered in eukaryotes in 1978, when Walter Gilbert proposed that variations in splicing sites could expand the coding capacity of genes (Gilbert 1978). Alternative splicing is widely prevalent within organisms, and in humans, it has been discovered that 92-94% of multi-exon genes undergo alternative splicing events (Wang et al 2008).

Prokaryotic genes typically possess a polycistronic structure, containing multiple open reading frames (ORFs), with different ORFs encoding proteins of distinct functions. In contrast, eukaryotic genes are generally monocistronic, where each gene typically produces mRNA containing a single ORF. However, eukaryotic genes can encode multiple protein isoforms through the mechanism of alternative splicing. The splicing process is coordinated by the spliceosome, a large protein complex composed of multiple small nuclear ribonucleoproteins (snRNPs) and associated proteins(Matera and Wang 2014). This complex recognizes consensus sequences at the 5’ and 3’ ends of introns in pre-mRNA and accomplishes splicing through two transesterification reactions: the first cleavage occurs at the 5’ GU splice site, and the second at the 3’ AG splice site, subsequently ligating the exons(Marasco and Kornblihtt 2023).

Alternative splicing events are classified into seven types: Alternative First Exon (AF), Alternative Last Exon (AL), Alternative 5’ Splice-Site (A5), Alternative 3’ Splice-Site (A3), Retained Intron (RI), Skipping Exon (SE), and Mutually Exclusive Exons (MX). Among these, Skipping Exon events is the most prevalent in animals(Keren et al 2010), while Retained Intron events are most common in higher plants. Specifically, in *Gossypium raimondii*, retained intron events constitute 40% of the identified alternative splicing events(Li et al 2014).

Transposable elements (TEs) are sequences capable of dynamic repositioning within the genome, first discovered by Barbara McClintock in maize at the Cold Spring Harbor Laboratory(Mcclintock 1950). TEs can be divided into two main classes: Class I and Class II. Class I, also known as retrotransposons, operate on a "copy-and-paste" mechanism and include Long Terminal Repeats (LTRs), Long Interspersed Nuclear Elements (LINEs), and Short Interspersed Nuclear Elements (SINEs)(Finnegan 1989). LTRs are commonly found in plants, while LINEs and SINEs are more prevalent in animals(Vitte and Panaud 2005). On the other hand, Class II TEs, also known as DNA transposons, function through a "cut-and-paste" mechanism. And they are further categorized based on the presence of a Terminal Inverted Repeat (TIR) domain into subclass I and subclass II(Wicker et al 2007). In eukaryotes, the distribution of transposable elements (TEs) within the genome is not random. When integrating into the genome, TEs exhibit specific preferences for certain locations. Additionally, they undergo a selection process, where only insertions at certain positions are more likely to be preserved. This highlights the complex interaction between TEs and the genomic landscape in which they operate(Sultana et al 2017). The long terminal repeats (LTRs) of endogenous retroviruses (ERVs) provide regulatory templates for stage-specific transcription initiation during early human embryonic development, which results in the coordinated expression of hundreds of ERV-derived RNA transcripts, playing a crucial role in regulating embryogenesis(Göke et al 2015).

RNA sequencing technology has developed over several decades. Short-read sequencing, represented by Illumina and MGI, is widely used in transcriptomic studies due to its high throughput and low cost. However, due to the limitations in read length, it cannot capture the complete mRNA sequences, and the accuracy of algorithmically assembled short read fragments is not guaranteed(Stark et al 2019). In contrast, long-read RNA sequencing approaches are capable of systematically deciphering the features of isoforms and fully identifying the prevalence of alternative splicing (AS) by generating sequences that span entire isoforms(Li et al 2017). Currently, long-read sequencing technology is dominated by a select few companies.

In 2011, Pacific Biosciences (PacBio) launched the Single-Molecule Real-Time (SMRT) full-length RNA sequencing technology, also known as Iso-seq. This technology enhances the accuracy of RNA-seq at the isoform level by providing high-resolution maps of complete cDNA sequences without the need for assembly. However, due to its initially high cost, it did not become widely used until 2015. The Circular Consensus Sequencing (CCS) mode produces high-quality, highly accurate consensus reads(Rhoads and Au 2015).

In 2014, Oxford Nanopore Technologies (ONT) released its first commercial product, the MinION. Since that time, the technology has advanced considerably, achieving substantial enhancements in accuracy. Presently, ONT employs two main sequencing strategies: the first is based on cDNA synthesis, which aligns with the principles used by most current sequencing technologies; the second strategy allows for the direct sequencing of single-stranded RNA without the necessity of reverse transcription, enabling sensitive detection of nucleotide modifications present on native RNA(Jain et al 2022).

In this context, it is crucial to differentiate between Differential Gene Expression (DGE), Differential isoform Expression (DIE), and Differential isoform Usage (DIU). DGE examines changes in gene-level expression across different conditions. For example, if gene A exhibits a change in overall expression between two conditions, it is considered differentially expressed. DIE, however, focuses on changes in the expression of individual isoforms—different isoforms produced by a gene. For instance, if two of the three isoforms of gene A (A.2 and A.3) show changes in their expression levels, which is noted under DIE. In Differential Isoform Usage (DIU), shifts are observed in the expression of different isoforms of the same gene across various developmental stages. For example, even if the total expression levels of Gene A and Gene B remain consistent, isoform switching can occur, which refers to changes in the dominant isoform. DIU focuses on changes in the proportions of various isoforms within the total gene expression. For instance, in condition 1, isoform A.2 might represent 60% of the total expression, with A.1 and A.3 each contributing 20%. In condition 2, although the total expression of gene A remains nearly unchanged, the proportion of A.3 could increase to 50%, while A.1 and A.2 account for 25% each, thereby indicating a significant shift in isoform usage(Froussios et al 2019). Additionally, research indicates that isoforms from the same gene can have different functions(Baralle and Giudice 2017). Therefore, limiting analysis to gene-level differential expression may not yield a comprehensive or thorough understanding, underscoring the potential importance of DIE studies over DGE. In various conditions, even if no significant gene-level differential expression is observed, significant DIE may still occur among the isoforms.

Exploring isoform diversity in *Gossypium barbadense* and *Gossypium hirsutum* based on the PacBio Iso-seq system. The allopolyploid *Gossypium barbadense* (3-79) and *Gossypium hirsutum* (TM-1) are the current main two cultivated varieties. Although the genome assembly level has been well established(Wang et al 2019), the gene structure annotation and isoform discovery are not comprehensive. Integrating full-length isoform structures identified based on SMRT-seq to improve the completeness and comprehensiveness of annotations, this research generates a transcriptome map with novel isoforms and more comprehensive details. Additionally, it explores isoform-resolved transcriptional changes during 10 developmental time points of cotton fiber based on high-depth RNA-seq, identifying key isoforms related to fiber development.

Further explore dynamics of differential isoform expression and alternative splicing over fiber developmental stage transitions.

## Result

At the National Center for Biotechnology Information (NCBI), we integrated data from multiple projects. Specifically, from project PRJNA726938, we obtained 4,022,742 full-length non-concatemer (FLNC) reads for the root, leaf, petal, and stem of *Gossypium barbadense* (3-79) and 3,657,099 FLNC reads for *Gossypium hirsutum* (TM-1), both generated on the PacBio Sequel II platform. Additionally, from project PRJNA359724, we collected 517,229 FLNC reads from *Gossypium barbadense* fiber at developmental stages 0 DPA, 7 DPA, 10 DPA, 12 DPA, 25 DPA, and 30 DPA, generated on the PacBio RS II platform. Furthermore, from project PRJNA503814, we obtained 415,159 FLNC reads from root, leaf, flower, stem, ovules at 0 DPA, 5 DPA, 10 DPA, 20 DPA, and fiber at 10 DPA and 20 DPA of *Gossypium hirsutum*, also generated on the PacBio RS II platform. Through these efforts, we constructed the most complete and comprehensive reference transcriptome for cotton to date, identifying 183,767 isoforms for *Gossypium barbadense* (3-79) and 178,994 isoforms for *Gossypium hirsutum* (TM-1).

In the China National GeneBank DataBase (CNGBdb) under accession CNP0002656(Zhang et al 2022), RNA sequencing data were collected from ten time points during fiber development in *Gossypium barbadense* (3-79) and *Gossypium hirsutum* (TM-1). The time points include -3 DPA, -1 DPA, 0 DPA, 1 DPA, 3 DPA, and 5 DPA for ovule development, as well as 10 DPA, 15 DPA, 20 DPA, and 25 DPA for fiber development.

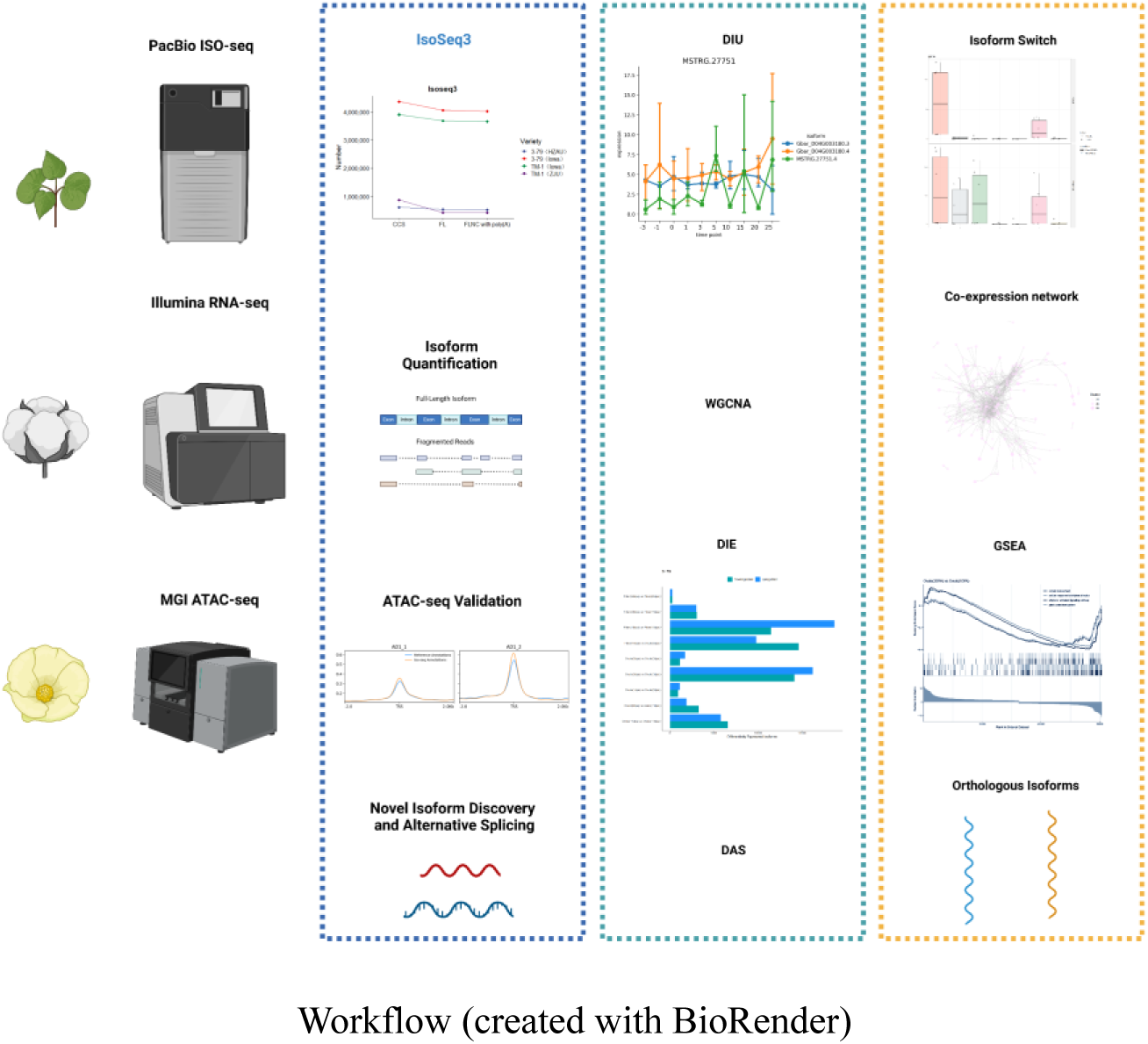

The cotton industry holds a critical position in the national economy, and enhancing the quality of cotton fibers remains a key objective for crop breeders. The quality of these fibers is intimately linked to their developmental processes. Despite significant progress, the understanding of the molecular mechanisms occurring during fiber development to further to explore.

High-throughput next generation sequencing has expanded the knowledge of the cotton transcriptome, facilitating gene expression profiling on a massive scale. Nevertheless, current cotton gene annotation mainly derives from short-read RNA-seq. Limited to read length, such technologies are inherently difficult to capture the contiguous sequence of mRNAs, frequently inducing to fragmentation, incompletion, or compression error in isoform annotations. The advent of PacBio SMRT-seq overcomes these limitations, with read lengths reaching up to 25kb(Rhoads and Au 2015), eliminating the need for assembly and fundamentally eradicating transcript assembly errors. Iso-seq has been widely applied to maize(Wang et al 2016), sorghum(Abdel-Ghany et al 2016), human and mouse cerebral cortex(Leung et al 2021), and *Fusarium graminearum*(Lu et al 2022).

The current PacBio iso-seq unable to be straightforwardly used for quantification, therefore, it is necessary to integrate additional short-read RNA-seq to improve the precision of isoform quantification(Amarasinghe et al 2020). Relying solely on short-read RNA-seq can miss unannotated isoforms, leading to an incomplete analysis of RNA expression and splicing during cotton fiber development stages. In fact, alternative splicing (AS) has been proven to be crucial for the development of cotton fibers(Zheng et al 2023).

Integrating full-length isoform charactered by Iso-seq with the high read-depth of RNA-seq, construct the most comprehensive isoform-resolved catalog of transcriptional changes across time points in fiber development.

**Fig 1.**
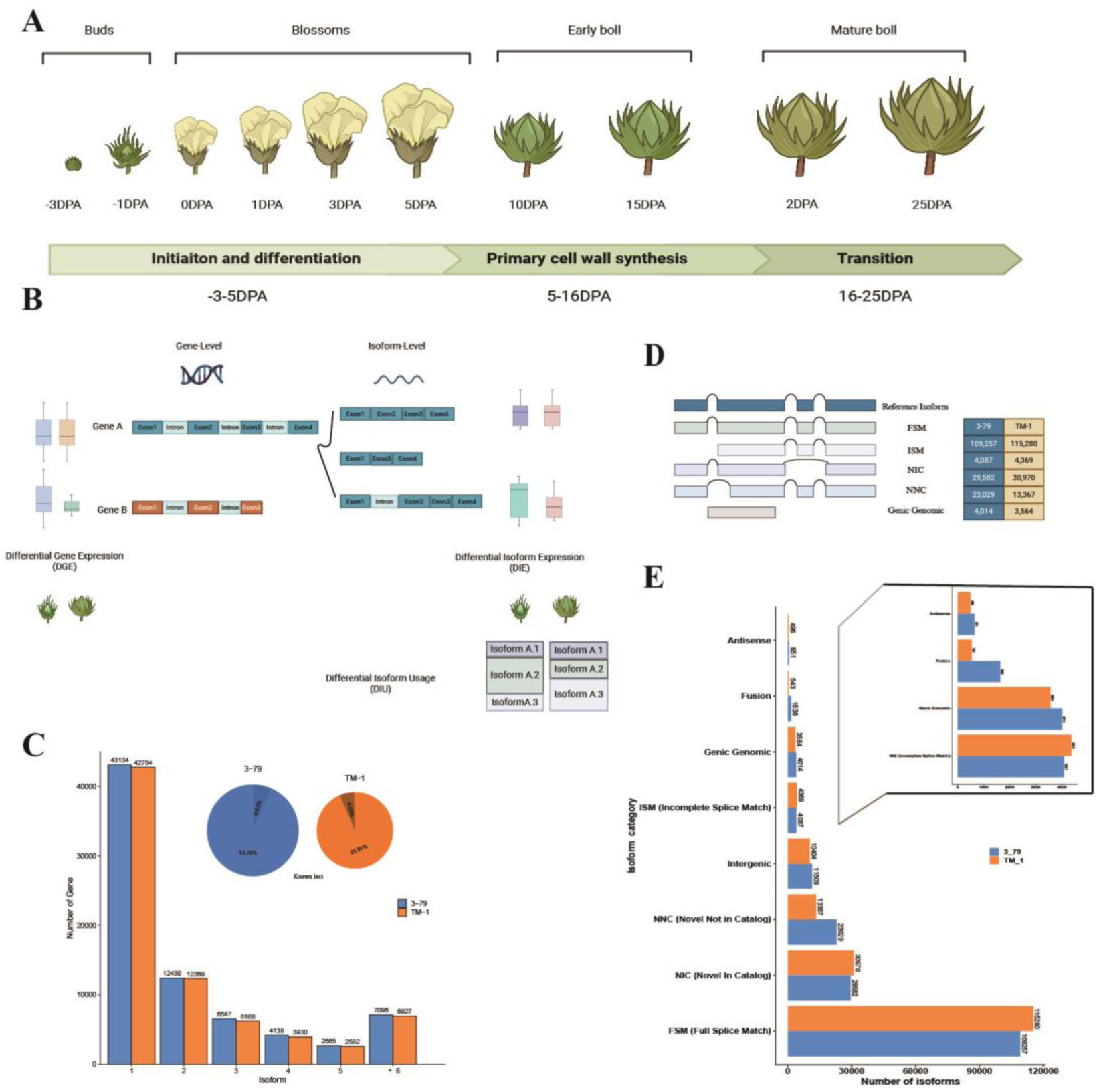
Generation and basic statistical analysis of the isoform-resolved cotton transcriptome. **A** Overview of Cotton Fiber development stages (created with BioRender). **B** The distinctions between differential gene expression (DGE) and differential isoform expression (DIE) and what Differential Isoform Usage (DIU) entails (created with BioRender). **C** The number of isoforms identified per gene in 3-79 and TM-1 and isoforms are annotated to regions on the reference genome. **D** The primary structural categories for isoforms models for genes. **E** Number of isoforms in the *Gossypium barbadense* (3-79) and *Gossypium hirsutum* (TM-1) transcriptome for each structural category.

In the constructed novel reference transcriptome, isoforms can be categorized based on whether their splice sites (SJs) match known isoforms into several types: Full Splice Match (FSM), Incomplete Splice Match (ISM), Novel In Catalog (NIC), Novel Not in Catalog (NNC), Fusion, Genic Genomic, Intergenic, and Antisense. Among these, FSM constitutes the majority in samples 3-79 (59.45%) and TM-1 (64.40%).

Both FSM and ISM splice junctions match annotated isoforms, indicating that they are transcribed from known genes into known isoforms. However, a significant portion of the isoforms are not present in the existing annotations and are referred to as Novel Isoforms. NIC and NNC are new isoforms transcribed from known genes. Antisense and Intergenic do not match genomic annotations and belong to novel genes, meaning they are transcribed from novel genes into novel isoforms. Fusion occurs when multiple known isoforms merge together. Genic Genomic lies within known gene regions but does not fully match any known isoform models and can be understood as isoform variants of known genes.

In eukaryotic cells, nuclear DNA is not found in a naked state but is instead bound to histones. DNA wraps around these histones to form structures known as nucleosomes, which resemble beads. These nucleosomes are further folded and aggregated to form the condensed structures known as chromosomes(Richmond and Davey 2003). Transcription requires unraveling the higher-order structure of DNA, but only opens the region where the gene is expressed(Li et al 2007). The utility of Tn5 transposase in the selective cleavage of unprotected DNA regions by proteins (Buenrostro et al 2013). The Assay for Transposase-Accessible Chromatin with high-throughput sequencing (ATAC-seq) is a classic method for assessing chromatin accessibility. By collecting existing ATAC-seq data, it was found that humans and mice account for 90% of the data, while other species are still in the process of accumulating data (Luo et al 2022). Presently, ATAC-seq data is unavailable for Gossypium barbadense (3-79) in cotton research, and the data for TM-1 is limited solely to leaf samples. DNase-seq is predicated on the sensitivity of active transcriptional regions, such as promoters and introns of expressed genes, to DNA endonucleases, indicating that the chromatin in these regions is in an accessible state (Minnoye et al 2021). The publicly available data includes leaf DNase-seq data for the 3-79. According to Supplemental Fig S1, ATAC-seq and DNase-seq reads are expected to be enriched around the nucleosome-free regions of transcription start sites (TSSs). In two biological replicates of 3-79 and TM-1, we observed greater enrichment at TSSs identified by Iso-seq, suggesting that the TSSs annotated by Iso-seq are more accurate.

**Fig. 2.**
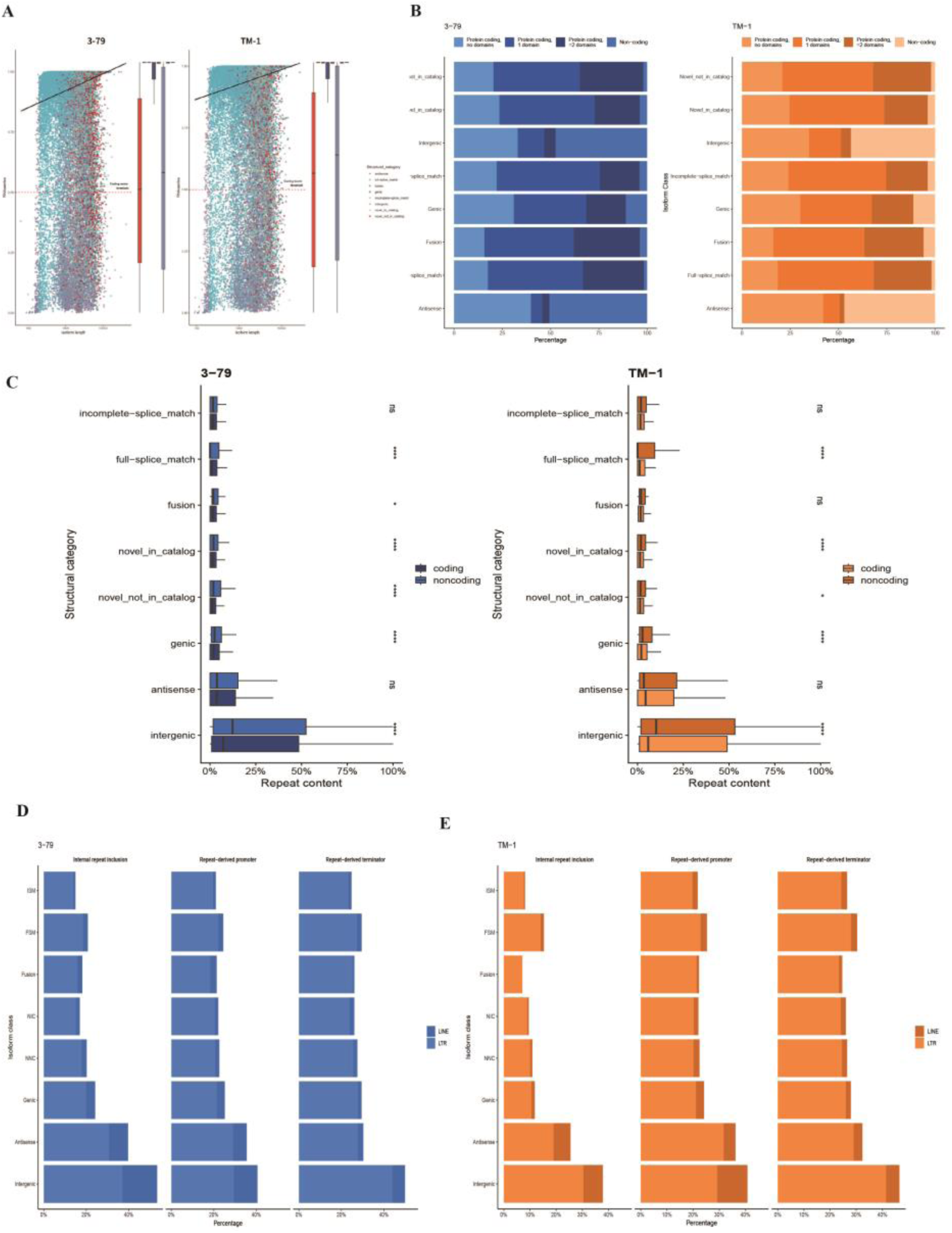
Comprehensive functional analysis of isoform-resolved transcription in cotton. **A** Scatter plot displaying isoform length and based on RNAsamba predicting coding potential for each isoform, colored by isoform categories. Boxplots display the distributions of isoform coding potential along Y axes. **B** Bar plot illustrating the distribution of isoforms in each category on their predicted protein-coding status and the presence of known protein domains in the encoded peptide both 3-79 and TM-1. **C** Box plots illustrating the relative repeat content of isoforms in each structural category by predicted protein coding status. **D** Bar plots illustrating the number of repetitive elements originating from promoters, internal regions, and terminators for each isoform category, classified by repeat class in 3-79. **E** Bar plots illustrating the number of repetitive elements originating from promoters, internal regions, and terminators for each isoform category, classified by repeat class in TM-1.

According to RNAsamba(Camargo et al 2020), the protein-coding potential of each isoform has been assessed. It was found that coding potential positively correlates with isoform length, and isoforms transcribed from known genes exhibit significantly higher coding potentials compared to those from novel genes. The relationship between coding ability and length for isoforms in species 3-79 and TM-1 appears to be essentially similar. Based on the standard set by RNAsamba, values greater than 0.5 typically indicate coding capability, with higher values suggesting stronger coding potential. The relationship between isoform length and coding ability is not straightforwardly linear and cannot be fully explained by conventional linear models. The analysis indicated that the coding potential of an isoform generally increases with its length, likely due to previous incompleteness of isoform measurements. As the completeness of isoforms improves, some that were previously not predicted to code begin to show coding potential due to increased completeness. This underscores the importance of enhancing the examination of isoform completeness.

First, determine whether isoforms are coding using RNAsamba(Camargo et al 2020), a deep learning-based identification method. If they are coding, identify protein domains from the predicted coding isoform by applying InterProScan(Jones et al 2014). InterPro is a comprehensive database that integrates multiple protein signature databases, including Pfam(Mistry et al 2021), PRINTS, ProDom, SMART, TIGRFAMs, and PANTHER. Pfam, particularly noted for its focus on predicting protein domains, is utilized to identify domains in coding isoforms.

Isoforms from novel genes are largely predicted to be non-coding or coding but without annotated known domains. In contrast, isoforms from known genes are primarily predicted to produce proteins that contain at least one recognized domain. Fusion isoforms, which result from the merging of multiple known isoforms, might involve combinations of functional regions from several proteins. Furthermore, genic regions, due to their inclusion of more genomic sequences such as introns and non-coding regions, might retain potential domains that would not normally be translated. The potential inclusion of Intron 1 in the final mRNA transcript can result in the introduction of unique sequences that are typically absent from proteins. If these intron-derived sequences possess specific structural or functional motifs, such as the potential to form a distinct domain, the resulting protein product may exhibit novel functional characteristics. The incorporation of these atypical sequences through the retention of the intron can confer unique properties to the protein, potentially expanding its functional repertoire or altering its characteristic.

To elucidate the role of Transposable element in the transcriptome, RepeatMasker(Smit et al 2015) was employed to identify repetitive elements within isoforms. FSM and ISM, also referred to as known isoforms, exhibit the lowest content of repetitive sequences, which is almost negligible. Additionally, it is apparent that noncoding isoforms consistently harbor a substantially greater proportion of repetitive content in comparison to their protein-coding counterparts. The same trend has been observed in other isoform types, particularly antisense and intergenic isoforms, which also arise from novel gene loci. This is likely because most isoforms in these categories do not encode proteins. Antisense and intergenic isoforms are implicated in gene regulation, potentially through the involvement of repeat sequences. For a more detailed investigation, repeat sequences were categorized and extracted from specific genomic regions: 2kb upstream of the Transcription Start Sites (TSS) for promoter regions, the isoform regions themselves, and 2kb downstream of the Transcription End Sites (TES) for terminator regions.

In the identified transposable elements, LTR and LINE are the predominant types, while other transposon types are exceedingly rare. LTRs are the principal components, primarily distributed across FSM and intergenic isoforms. FSM isoforms, which match perfectly with known splice junctions and are evolutionarily conserved, are likely to contain numerous known TEs, potentially influencing gene expression regulation. Conversely, the underrepresentation of TEs in ISM isoforms may stem from incomplete splicing sites, incorporating only fragments of exons and splice sites, thus posing challenges in TE identification. Intergenic areas, possibly subjected to lower selective pressures, may allow more transposon activity and accumulate a greater number of TEs. The distribution of LTR-associated promoters and terminators on intergenic and antisense isoforms shows similarities to that on other isoforms. However, significant variations are observed in FSMs, ISMs, NIC, and NNC isoforms. This variability suggests that transposons in regions upstream of transcription start sites and near transcription termination sites may facilitate the generation of diverse isoforms variants through mechanisms like alternative initiation and termination, which might evade detection in the main isoform regions.

**Fig. 3.**
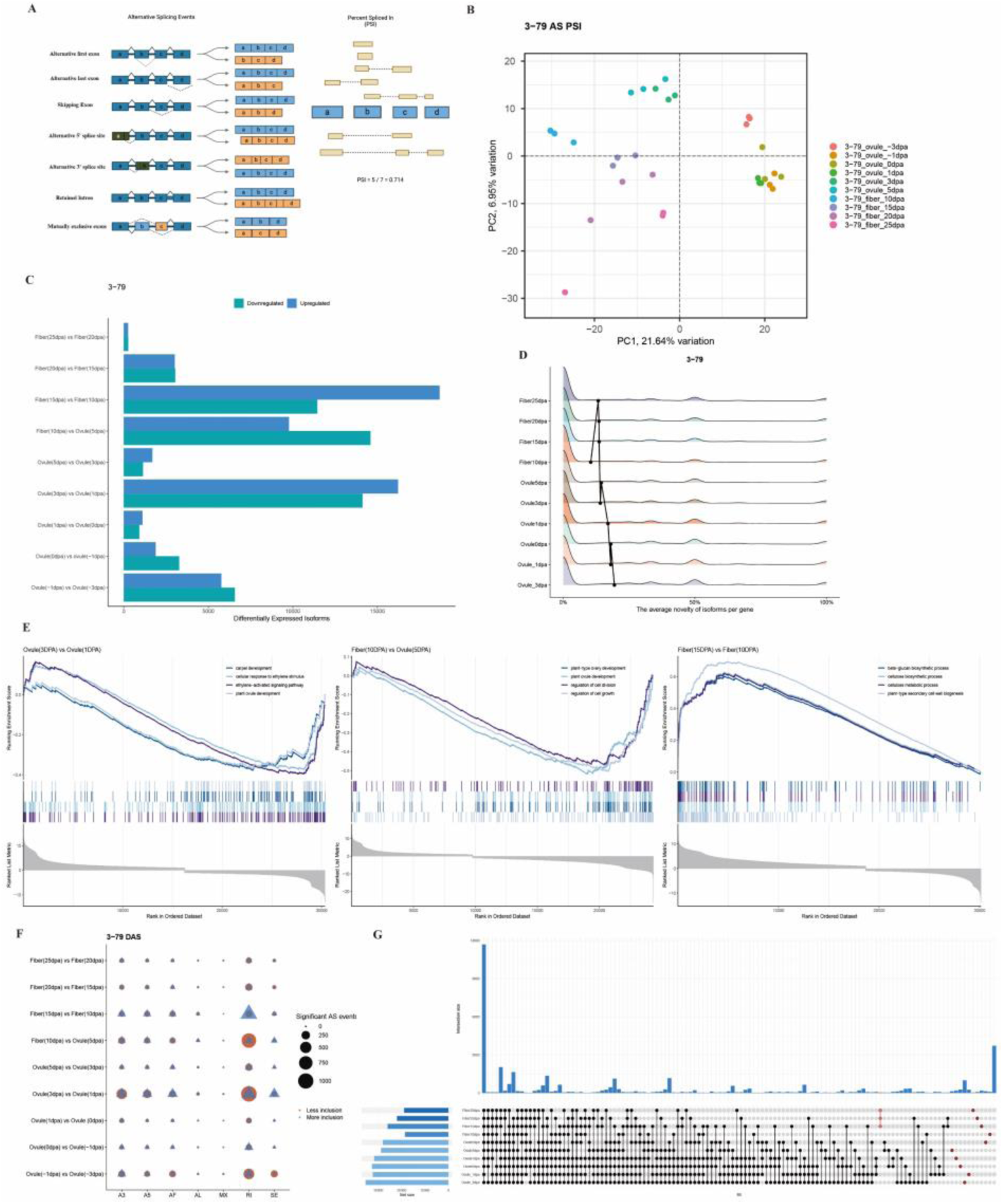
Exploring the impact of isoform diversity on cotton fiber development: Insights and Implications in 3-79. **A** Schematic representation of the seven AS events and the Percent Spliced In (PSI) values quantify alternative splicing events by calculating the ratios of junctions detected to annotated splice junctions (created with BioRender). **B** Performing PCA on alternative splicing PSI values from both ISO-seq and RNA-seq data, with points colored according to developmental stages. **C** Number of significantly differentially expressed isoforms at each fiber development stages transition in *Gossypium barbadense*. **D** The percentage of short RNA-Seq reads mapped to novel isoforms of each gene expressed at each developmental stage. **E** GSEA of isoform-level differential expression at three critical time points during fiber development. **F** Number of significant AS events at each fiber development stages transition in *Gossypium barbadense*. **G** Upset plot of Retained intron events characteristic of fiber development stages.

Based on the inclusion or exclusion of exons, alternative splicing events can be classified into seven types: Alternative First Exon (AF), Alternative Last Exon (AL), Skipping Exon (SE), Alternative 5’ Splice Sites (A5), Alternative 3’ Splice Sites (A3), Retained Intron (RI), and Mutually Exclusive Exons (MX). The quantitative assessment of alternative splicing events employs the Percent Spliced In (PSI) metric, which quantifies the inclusion of a specific exon within an isoform. The calculation formula for PSI is as follows: 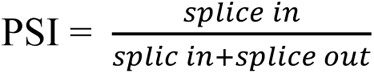, where ’splice in’ refers to the number of reads that include the exon in the final mRNA as part of a specific alternative splicing event, and ’splice out’ refers to the number of reads where the exon is excluded from the final mRNA. Thus, PSI represents a ratio that reflects the relative frequency of a specific splicing event, aiding in understanding how splicing patterns vary across different developmental stages. However, it’s important to note that this metric quantifies the overall mRNA levels in a mixed-state sample, rather than focusing on the alternative splicing of any specific mRNA.

Dynamic changes in PSI of gene alternative splicing events have been further explored in the fiber development process. A Principal Component Analysis (PCA) of PSI across isoforms identified from Iso-seq and quantified by RNA-seq revealed distinct separations between stages: from -3 DPA ovule to 1 DPA ovule, from 3 DPA ovule to 5 DPA fiber, and from 10DPA fiber to 25 DPA fiber. These findings corroborate the general trends previously described in the literature on fiber development. Prior research delineated the stages of fiber development, identifying the period from -3 DPA to -5 DPA as the initiation phase, 10 DPA and 15 DPA as the primary cell wall synthesis phase, and 20 DPA to 25 DPA as the transition from the primary to the secondary cell walls. However, some distinct findings indicate that 1 DPA may also mark a critical juncture. This observation corresponds with studies indicating that the interval from -3 DPA to 3 DPA represents the initiation phase of cotton fiber development, characterized by the protrusion and enlargement of trichomes on the epidermal surface(Qin and Zhu 2011). Importantly, the stage at 1 DPA ovule is pivotal for determining the number of fiber cells, thereby constituting one of the critical developmental periods that influence the yield traits of cotton(Prakash et al 2020). This suggests that the initiation phase can be subdivided further into two distinct stages, with 1 DPA acting as a subtle, yet pivotal, transition point. This novel insight is fascinating and deserves further exploration.

To assess the extent to which novel isoforms contribute to gene expression levels throughout developmental stages. Initially, RNA-seq data was used to identify reliably expressed isoforms at various stages of fiber development, followed by calculating the percentage of novel isoforms arising from individual genes. The highest levels of isoform novelty were observed in the ovules at -3 DPA and -1 DPA, where many unannotated isoforms were identified. This novelty decreased progressively through the developmental stages, reaching its lowest at 10 DPA fiber. This trend could be attributed to incomplete annotations in the earlier stages, such as at -3 DPA and -1 DPA, while by 10 DPA, the involved isoforms were better recognized and described within the existing genomic annotations. Additionally, the complexity of transcription mechanisms in the early stages of fiber cell development might contribute to this pattern. However, as development progresses, the focus of fiber cells shifts towards cellulose synthesis and cell wall fortification, stabilizing the gene expression pattern.

To assess isoform-level differential expression, a differential expression analysis was conducted using DESeq2 ((Love et al 2014). The results revealed that isoforms showed the most significant differences during the transition from 10DPA to 15DPA, followed by transitions from 1DPA to 3DPA and 5DPA to 10DPA. These notable differences at specific developmental transitions merit further investigation. Enrichment analysis was conducted on these significantly different isoforms to explore their potential functions.

GSEA (Gene Set Enrichment Analysis) offers a more inclusive approach compared to GO (Gene Ontology) analysis by evaluating all genes without the limitations imposed by individual gene significance thresholds. By ranking genes based on their expression changes (Fold Change), GSEA assesses the enrichment of specific gene sets within the complete gene ranking, which effectively circumvents the risk of excluding critical genes due to stringent criteria (Subramanian et al 2005). The top section of the diagram features a line chart representing the Enrichment Score (ES). The middle section, termed the Hits chart, visually resembles a barcode and marks the positions of each gene. The lower section illustrates the distribution of rank metric values for all isoforms, sorted in ascending order. The peak’s left side identifies the core genes. An ES value above zero suggests that the gene set is upregulated, whereas a value below zero indicates downregulation.

During the 1DPA to 3DPA period in cotton fiber development, there is a noted downregulation in the gene sets expression associated with ovule development, ethylene-activated signaling pathways, cellular responses to ethylene, and carpel development. Plant hormones are pivotal in fiber development and elongation, with various hormones synergistically contributing to the growth process(Ahmed et al 2018). The GhXB38D regulates ethylene biosynthesis during cotton fiber elongation. Research indicates that inhibiting GhXB38D expression during the rapid elongation phase significantly enhances fiber length(Song et al 2023). Furthermore, increasing cytokinin levels and suppressing GhCKX expression can augment cotton fiber yield(Zhao et al 2015). The cellular response to ethylene likely modulates key mechanisms governing cell division and elongation. Ovule development provides the essential structural and physiological foundation for fiber initiation and growth, with fibers originating from the outer integument cells of the ovule. In this phase, the reduction in ethylene responsiveness leads to diminished signaling capabilities, initially manifesting as a deceleration in carpel development.

During the development of cotton fibers from 5DPA to 10DPA, gene sets associated with cell growth regulation, cell division regulation, plant ovule development, and plant-type ovary development exhibit notable downregulation. In this developmental stage, there is a deceleration in cell growth and a reduction in cellular differentiation. This transition is intricately linked to the shift in fiber cells from the division phase to the expansion phase. The growth of fiber cells requires meticulous regulation to ensure the appropriate cell size and morphology, as the cells begin to transition toward cell wall synthesis during this period.

During the transition from 10DPA to 15DPA, gene sets associated with four key pathways in cotton fibers exhibit upregulation. The four gene sets predominantly involve processes related to plant-type secondary cell wall biogenesis, as well as the metabolic and biosynthetic pathways of cellulose and beta-glucan. Commencing at 15DPA, the developmental focus of cotton fibers shifts from the synthesis of primary cell walls to that of secondary cell walls. Secondary cell walls, characteristically more robust and thicker than their Primary cell walls, are enriched with cellulose, lignin, and hemicellulose. While the synthesis of secondary cell walls becomes more prominent in subsequent developmental phases, early indicators of this process are discernible during the 10DPA to 15DPA period.

Differential alternative splicing analysis conducted using SUPPA2(Trincado et al 2018) revealed that retained intron is the most prevalent type of alternative splicing event in flower plants. Among all identified transcriptomic alternative splicing events, retained intron events not only represent the most numerous but also exhibit the most significant variations across different developmental stages. Specifically, during the development of cotton fibers from 10 DPA to 15 DPA, there is a greater retention of introns in isoforms at 15 DPA compared to 10 DPA. In contrast, there is less intron retention in isoforms at 10 DPA compared to 5 DPA, and at 3 DPA compared to 1 DPA. Therefore, it is essential to study the variations in intron retention events during the development of cotton fibers.

In the study of retained intron events specific to the developmental stages of cotton fibers, it has been observed that some genes exhibit RI only during certain periods, with the highest number of RI events occurring in the ovules at -3DPA. Besides these stage-specific intron retention events are predominantly concentrated in the period before flowering, from -3 to 0DPA.

Alternative splicing leads to variations in the expression levels of different isoforms of the same gene under different conditions. For example, two isoforms of a gene may undergo isoform switching(IS), exhibiting changes in their relative abundances across different time points. When two isoforms perform different functions, IS will affect the change in function(Lio et al 2024).

**Fig. 4.**
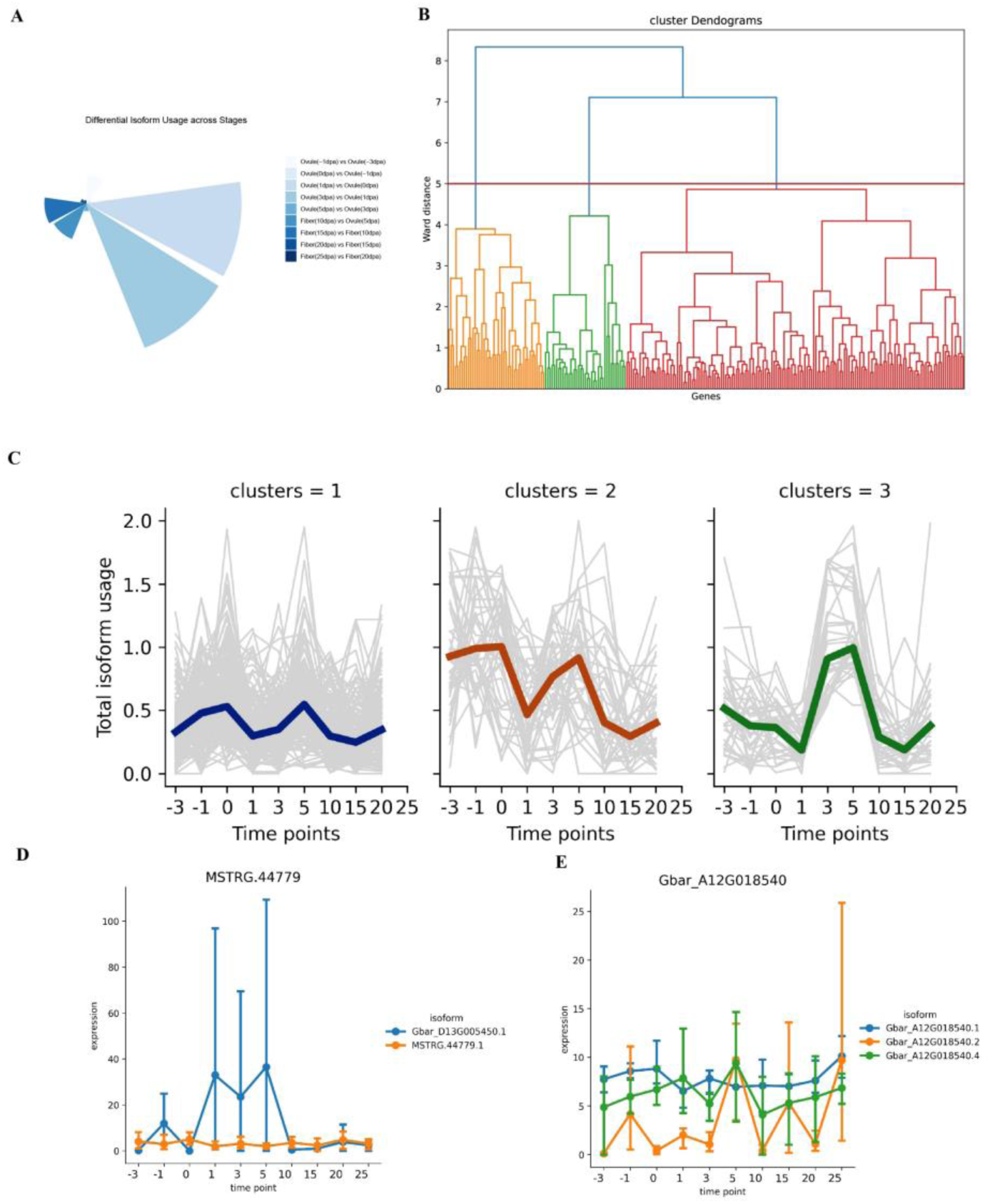
Systematic dynamic analysis of isoform switching during the time course of cotton development. **A** Number of isoforms with differential relative abundance at each fiber development stages transition in *Gossypium barbadense*. **B** Cluster dendrogram of hierarchical clustering 3-79 dataset. Each cluster is colored with different colors under the ward distance threshold at 5. **C** Cluster prototypes reveal the temporal pattern of total isoform usage changes across 10 time points. **D** Real examples of the changes in dominant isoforms at different time points.

Isoform usage measures the relative abundance of each isoform. Identifying isoform-switching behavior during fiber development can reveal critical changes at key time points. Using hierarchical clustering, genes with similar temporal expression patterns were grouped together. The total isoform usage changes for each gene were summed, displaying three distinct clustering patterns representing the trends in total isoform usage (the sum of the relative abundances of all isoforms). As a filtering criterion, we retained events with a relative abundance higher than 0.2 and an event importance score greater than 0.3. Genes undergoing isoform switching within the same cluster are expected to function synergistically with other genes related to specific biological processes. It can be observed that a significant number of isoform switching events occur at -1, 1, 3, and 5 DPA, while in TM-1, they primarily occur at -1, 0, 1, and 3 DPA. There are clear differences in isoform switching between the two species.

Weighted Gene Co-expression Network Analysis (WGCNA) is a scale-free network tool constructed based on systemic gene expression levels. It is used to explore whether genes exhibit similar trends of change across different treatments and to identify clusters of functionally similar genes. WGCNA was initially proposed by Professor *Peter Langfelder* from UCLA, and the corresponding analysis tool was developed in 2008 together with Horvath(Langfelder and Horvath 2008). It is important to note that some researchers have inappropriately used only a set of identified differentially expressed genes for co-expression network analysis in previous studies. This approach fails to consider potential interactions among genes within modules, resulting in a deterioration of the scale-free properties of the network when constructed solely with differentially expressed genes(Sánchez-Baizán et al 2022). When constructing an isoform expression network, the first step is to select the most appropriate β power to ensure the network meets the scale-free topology criterion. The type of network to be constructed should also be selected. Mean connectivity measures the degree of connection between an isoform and other isoforms, representing the average connectivity of all isoforms. A higher soft threshold (β power) brings the network closer to a scale-free topology, but it also decreases the Mean connectivity, so a balance must be considered.

**Fig. 5.**
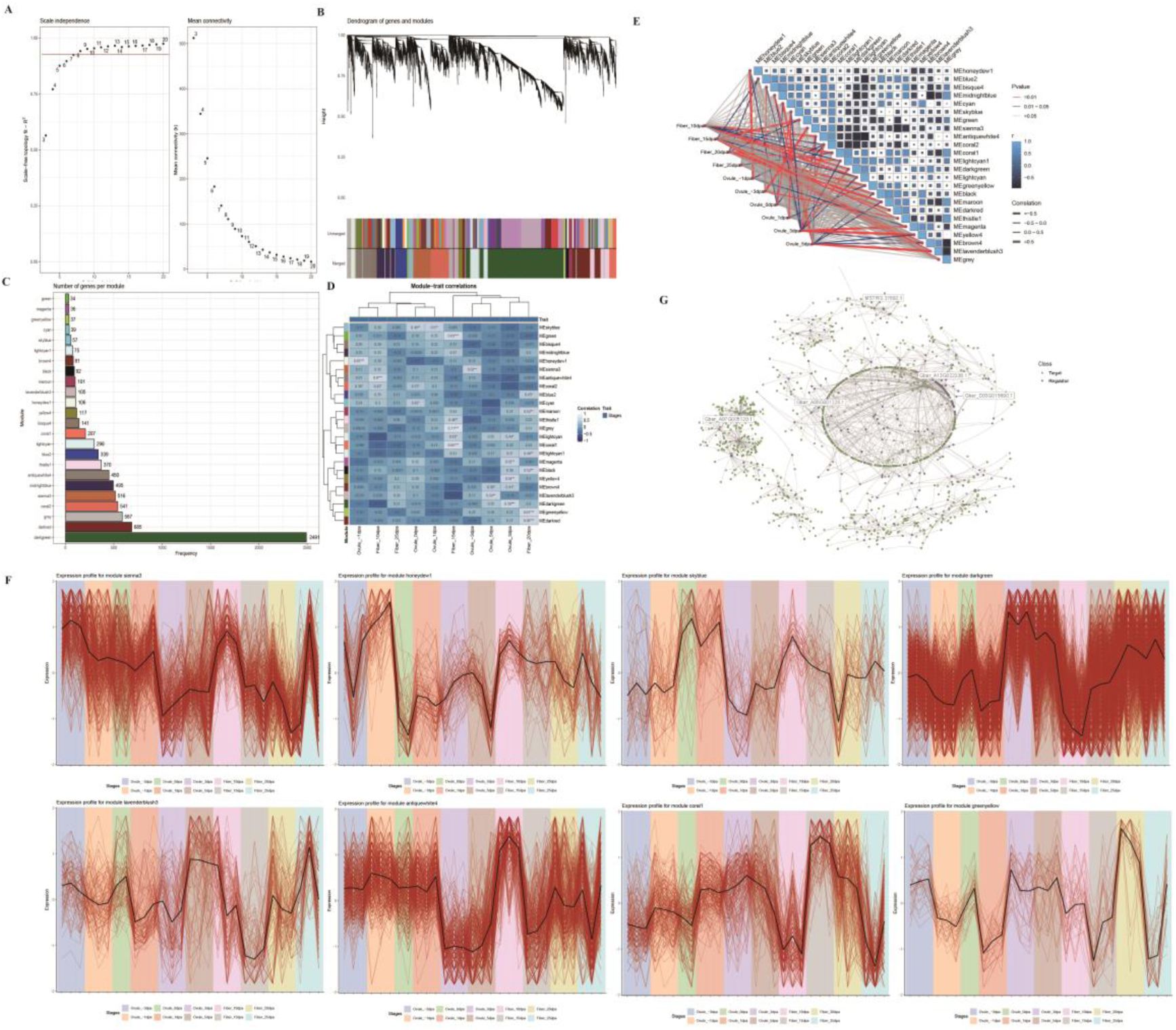
Co-expression network analysis across distinct developmental stages of cotton fiber. **A** Evaluation of the scale-free fit index and mean connectivity across a range of soft-thresholding power values. **B** Dendrogram of filtered isoforms clustered based on a dissimilarity measure (1-TOM) together with consistent module colors. **C** The number of isoforms per module. **D** Heatmap of the correlation between module eigengenes and traits of fiber developmental stages. Each cell displays the Pearson correlation coefficient and the corresponding P-value. **E** Overview of the correlation between isoform modules and specific stages of fiber development. **F** The expression profiles of the most significantly relevant periods corresponding to each module. **G** Isoform regulatory network inference.

This correlation heatmap clearly demonstrates the relationship between different stages of fiber development and their respective color-coded modules, which facilitates the identification of highly correlated isoforms within specific modules at particular stages.

Through co-expression network analysis, modules associated with different stages of fiber development are identified. In order to further narrow down the candidates to those isoforms that are most likely to explain stage-specific characteristics, a more reliable list of isoforms will be compiled using a machine learning approach for subsequent selection.

Regularized regression is extensively used for handling high-dimensional data, feature selection, and constructing predictive models, encompassing Ridge Regression, LASSO Regression, and Elastic Net. The Least Absolute Shrinkage and Selection Operator (LASSO), introduced by *Robert Tibshirani* in 1996(Tibshirani 1996), is a renowned method for variable selection. It regulates the estimated coefficients by adding a regularization term *λ* ∑^*p*^_j-1_ |*β*_*j*_|, to the objective function, thus *minimize*(*SSE* + *λ* ∑_j-1_^*p*^ |*β*_*j*_|) . In datasets with numerous features, LASSO effectively identifies and extracts the most crucial signal features. In medical research, employing the LASSO method to select disease-related genes from gene expression data effectively develops a robust prognostic gene signature associated with clear cell renal cell carcinoma(Yin et al 2021). LASSO regression is a powerful tool for feature selection, enabling the construction of a sparse model. Within the identified modules associated with fiber developmental stages, further refinement can be performed to isolate isoforms that are significantly linked to fiber development from a set of multiple predictor variables. LASSO is particularly advantageous in high-dimensional data settings, where isoforms are treated as predictor variables and fiber development stages as response variables. Building on the foundation of WGCNA, additional filtering is applied to the relevant modules for each developmental stage to enhance the selection of critical isoforms(Daneshafrooz et al 2022). In this approach, a specific module is selected for analysis, and variable selection is performed by shrinking certain coefficients to zero. Further analysis can then be conducted on the relevant modules corresponding to the 10 fiber developmental stages. Through the integration of WGCNA and LASSO, isoforms that are highly associated with fiber developmental stages can be effectively identified.

A coefficient profile plot against the log(λ) sequence in Lasso regression visually demonstrates how each predictor’s coefficient approaches zero as regularization strength increases, highlighting the variables that contribute most to the model at varying levels of penalty.

Figure 6B and Supplemental Figure S5 reveal significant differences in isoform expression during fiber development between sea island cotton and upland cotton. For example, the AP2/ERF domain, involved in ethylene signaling and fiber initiation in cotton, is expressed at -1 DPA in 3-79 and -3 DPA in TM-1, indicating that fiber initiation occurs earlier in TM-1. Phytocyanin proteins, commonly associated with photosynthesis, contribute to fiber development, with expression observed at 5 DPA in 3-79 and -1 DPA in TM-1, which suggests earlier activation in upland cotton.

**Fig. 6.**
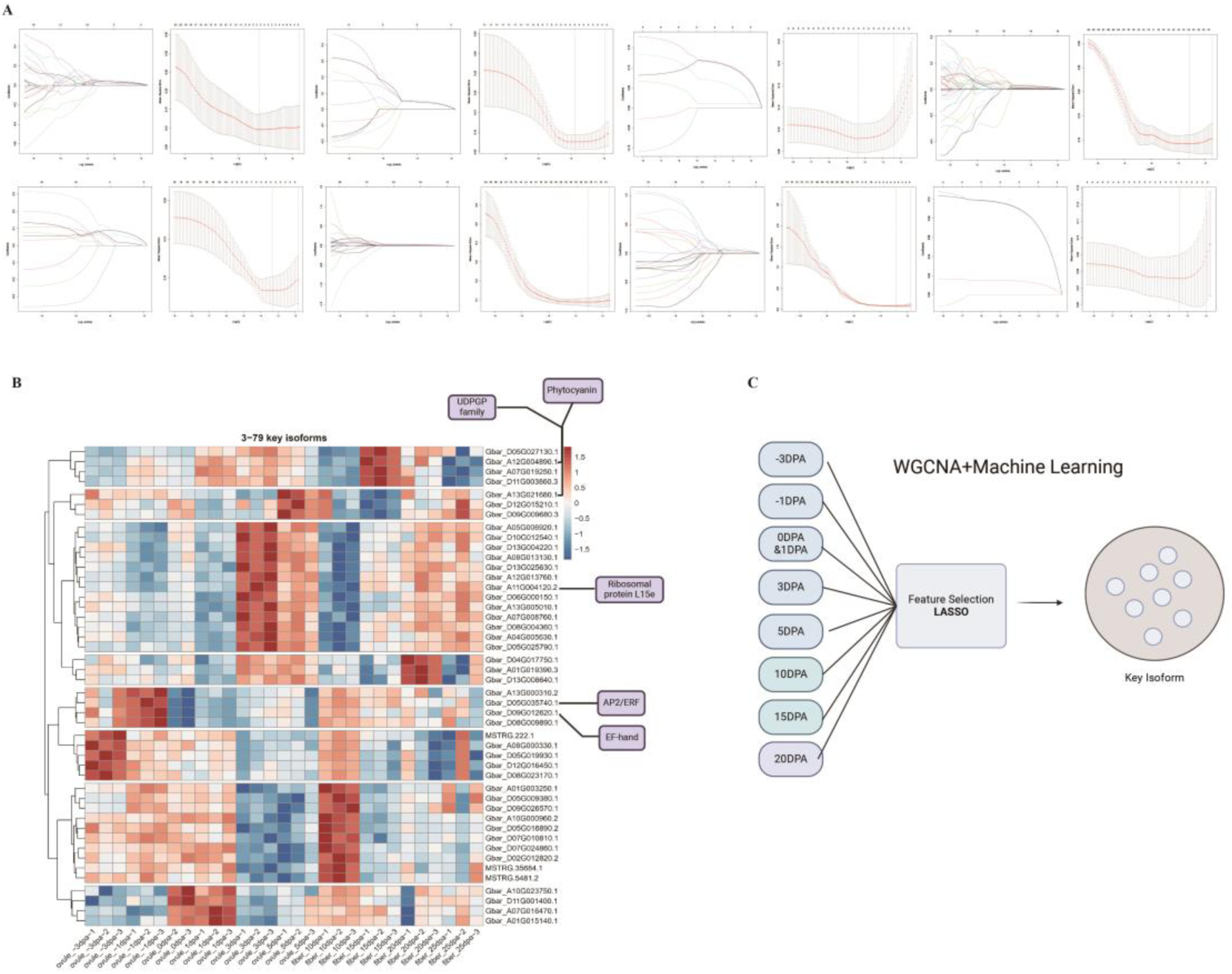
Integration of WGCNA with Machine Learning Techniques (LASSO). **A** Coefficient trajectory as log(λ) increases, showing variable significance under varying penalties. 5-fold cross validation for tuning parameter selection by LASSO regression. **B** Expression profile of identified key isoforms. **C** The schematic illustrates WGCNA combined with regularized LASSO regression analysis.

Ribosomal protein L15e, essential for protein synthesis, is particularly significant as fiber development is an energy-intensive process requiring substantial sugar metabolism for cellulose production. This protein is detected at 3 DPA in 3-79 and 1 DPA in TM-1, highlighting metabolic differences between the two species. Furthermore, UDPGP family enzymes, crucial for metabolic processes, are expressed at 10 DPA in sea island cotton and 5 DPA in upland cotton, suggesting that fiber elongation starts earlier in upland cotton. Lastly, the EF-hand domain, involved in calcium signaling, plays a key role in regulating cell wall expansion and cellulose synthesis during rapid fiber elongation. This domain is expressed at -1 DPA in sea island cotton and 5 DPA in upland cotton, again indicating that the developmental timeline is advanced in upland cotton compared to sea island cotton.

In summary, these findings suggest that fiber development in TM-1 occurs earlier than in 3-79, potentially leading to shorter fibers in TM-1 due to premature development, while the more gradual fiber elongation process in sea island cotton results in longer fibers.

Orthology refers to genes derived from a common ancestral species, subsequently passed down to each descendant species following speciation. In previous studies, orthologous groups were primarily identified at the gene level. However, due to the prevalence of alternative splicing, gene-level analysis alone does not fully capture the diversity involved in gene regulation. Therefore, extending the analysis to alternative splicing isoforms is essential. At the transcriptomic level, defining orthologous genes should account for alternative splicing. Based on the identification of orthologous isoforms, it becomes possible to compare differentially expressed isoforms and alternative splicing events between the two species, providing deeper insights into transcriptomic variation.

**Fig. 7.**
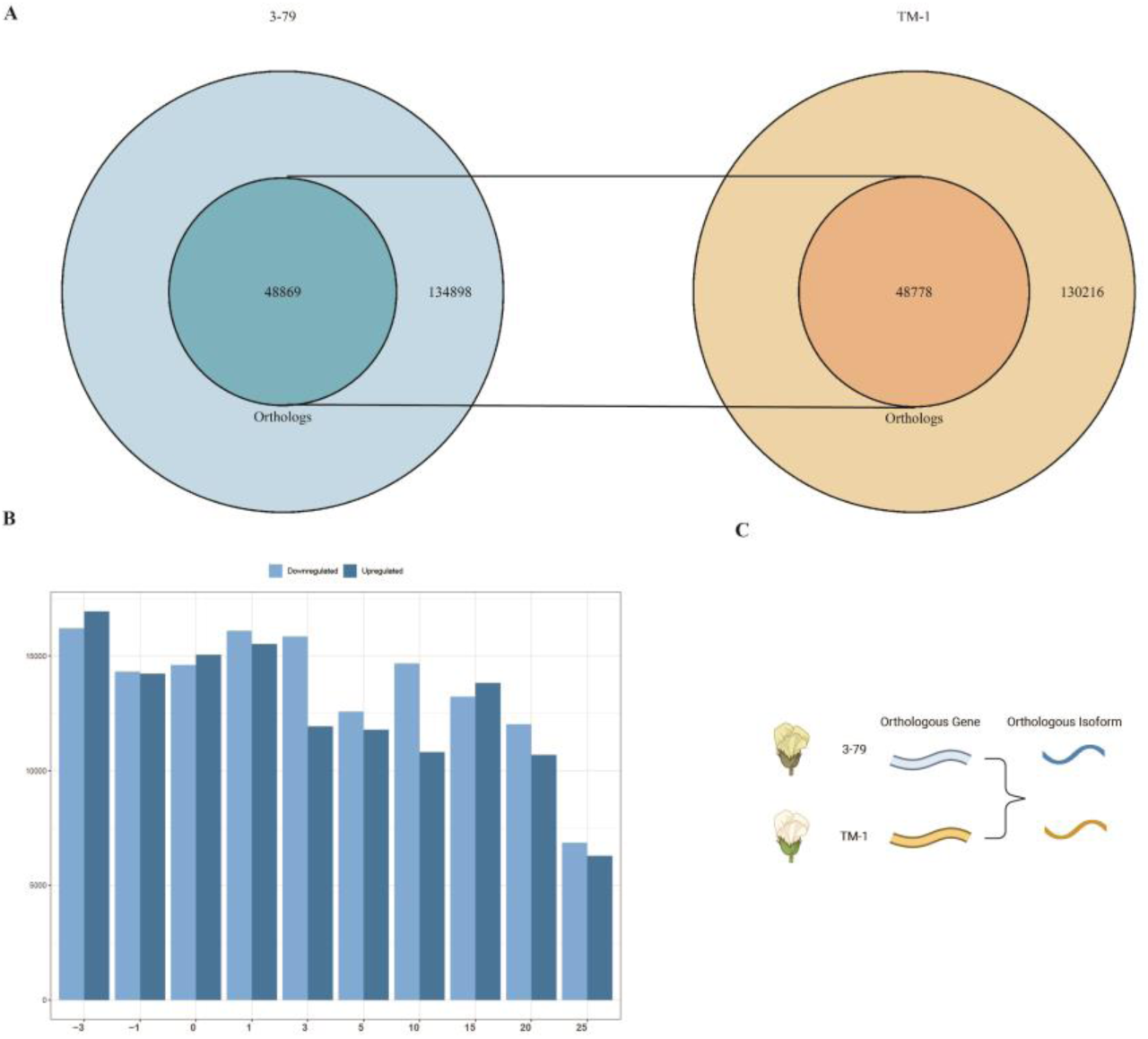
Cross-Species Transcriptomic Comparisons and Identification of Orthologous Isoforms. **A** Orthologous isoforms in 3-79 and TM-1. **B** Number of orthologous isoforms exhibiting significant differential expression in 3-79 and TM-1 at the same developmental time point. **C** Schematic diagram of identifying orthologous isoforms from orthologous genes.

Initially, 35,947 pairs of orthologous genes were identified using OrthoFinder(Emms and Kelly 2019). Within these, 28,290 pairs of orthologous genes were further analyzed, revealing 48,869 isoforms in 3-79 and 48,778 isoforms in TM-1 as orthologs. While nearly half of the orthologs are conserved between the two species at the gene level, notable differences persist at the isoform level, indicating substantial isoform diversity between 3-79 and TM-1. In both 3-79 and TM-1, the majority of isoforms are unique to each variety.

## Methods

### Construction of Full-Length Transcriptome Based on PacBio Iso-Seq

Raw sequencing reads were generated by PacBio platform and subsequently processed using the Iso-seq pipeline (3.8.2). Initially, subreads were processed using Circular Consensus sequencing (CCS) algorithm (6.4.0), applying a minimum of one pass to generate high-quality reads. Primers and SMRT adapters were then removed using Lima (2.7.1) to obtain full-length (FL) reads. Subsequently, concatemers were removed to produce Full-length Non-chimeric (FLNC) reads. Then FLNC reads were aligned to both the 3-79 (HAU.2) and TM-1 (HAU.1) genomes using Minimap2(2.17)(Li 2021). After alignment, transcript clustering was performed using TAMA(3.0)(Kuo et al 2020). Finally, isoform classification was carried out with SQANTI3(5.2.2)(Pardo-Palacios et al 2024).

### Analysis of publicly available ATAC-seq and DNase-seq data to validate Transcription Start Sites (TSSs)

The 3-79 leaf data was derived from publicly available DNase-seq, while the TM-1 leaf data was sourced from publicly available ATAC-seq. Quality control for the paired-end data was conducted using fastp (Chen et al 2018), followed by alignment with Bowtie2(Langmead et al 2019). The resulting BAM files were converted to BigWig format using the bamCoverage module from deeptools(Ramírez et al 2016). To identify chromatin accessibility, the intersection of the two biological replicates of the ATAC-seq and DNase-seq data was performed using Bedtools(Quinlan and Hall 2010). Subsequently, the chromatin accessibility at transcription start sites was examined across different isoforms, revealing accessible regions near the TSS.

### Integration of short-read RNA-seq data with long-read sequencing for full-length isoform quantification in the cotton fiber transcriptome

The obtained isoforms were compared and classified against known annotations, with splice sites validated using high-coverage short reads. Subsequently, false-positive isoforms were filtered using the machine learning algorithm sqanti3_filter.py, resulting in more comprehensive and accurate transcriptome. From the full-length transcriptome generated, isoforms across 10 developmental stages of cotton fiber were quantified using RNA-seq data. Salmon(1.10.2)(Patro et al 2017) was employed for isoform quantification, which, unlike traditional methods that rely on perfect alignment to the genome, leverages k-Mer matching between reads and isoforms. This pseudo-alignment approach offers superior efficiency and accuracy compared to traditional alignment-based methods. First, a transcriptome index was built, followed by isoform quantification, and the expression matrix from different developmental stages were merged. To streamline the quantification process, I developed a custom script based on Snakemake(5.4.0)(Köster and Rahmann 2012), which automated the entire workflow-from RNA-seq quantification to final expression matrix generation(https://github.com/Tang-pro/Snakemake_Isoquant).

### Analysis of protein-coding potential, protein domain content and repeat content

Open read frames (ORFs) were predicted from the nucleotide sequence of each isoform using RNAsamba(0.2.5)(Camargo et al 2020), with a scoring system applied. Isoforms with a coding potential score greater than 0.5 were classified as protein-coding. The predicted protein sequences were then scanned using InterProScan(5.55-88.0)(Paysan-Lafosse et al 2023), employing the Pfam database to identify the protein domains associated with each isoform. Additionally, repetitive sequences were identified using Repeatmasker(4.1.7)(Smit et al 2015) for both the transcriptome and the genome (2kb upstream of the transcription start site and 2kb downstream of the transcription termination site). For genome annotation, a species-specific repeat library was first constructed using RepeatModeler.

### Differential isoform expression analysis and alternative splicing analysis

After obtaining the isoform expression matrix in read counts, differential expression analysis was performed using DESeq2(1.44.0)(Love et al 2014). Differentially expressed isoforms were defined as those with an adjusted p-value ≤ 0.05 and log2FoldChange > 1 for upregulated isoforms or log2FoldChange < -1 for downregulated isoforms. Additionally, a custom script was used calculate the proportion of novel isoforms per gene across different developmental stages (https://github.com/Tang-pro/Novel_isoform_analysis_per_gene). Gene set enrichment analysis (GSEA) was conducted using clusterProfiler(4.12.6)(Wu et al 2021) in R 4.3.0. For alternative splicing event identification, SUPPA2(2.3)(Trincado et al 2018) was used. Seven types of alternative splicing events were identified based on the reconstructed GTF file: skipped exon (SE), alternative 5’ splice site (A5), alternative 3’ splice site (A3), retained intron (RI), mutually exclusive exons (MX), alternative first exon (AF), and alternative last exon (AL). Isoform expression data normalized to TPM, were combined with transcript TPM values from Salmon to calculate the Percent Spliced In (PSI). Differential splicing events were the identified using the SUPPA2 diffSplice module with a significance threshold of p-value ≤ 0.05 and differential PSI > 0.1 (more included) or PSI < 0.1 (less included). PSI values were further analyzed to detect splicing events specific to different stages, such as retained intron events. These events were visualized using Complexupset(1.3.3)(Lex et al 2014) to generate an upset plot. For differential isoform usage analysis, pairwise comparisons between different stages were conducted using IsoformSwitchAnalyzeR(2.4.0)(Vitting-Seerup and Sandelin 2019). However, since this method cannot handle comparisons across multiple time points, Spycone (Lio et al 2023) was used to identify isoform switch behaviors across 10 cotton fiber developmental stages.

### Weighted isoform co-expression network analysis combined with LASSO for further selection

Co-expression network analysis was performed on the normalized expression matrix using BioNERO(1.12.0)(Almeida-Silva and Venancio 2022). Initially, all isoforms with a median expression value below 10 were removed, followed by the identification of the most suitable β power that ensures the network satisfy the scale-free topology. Isoforms were then clustered, with modules of the same color representing group of isoforms with similar expression patterns. The co-expression network is represented as undirected weighted graphs, which does not reveal regulatory relationships. Therefore, to infer regulatory interactions, transcription factors were predicted from all protein sequences using PlantTFDB(Tian et al 2020), and the predicted structures were used to construct a regulatory network, illustrating interactions between regulators (transcription factors) and their targets isoforms. Although each module in the co-expression network contains a large number of isoforms corresponding to various fiber development stages, further filtering can be applied. Feature selection was conducted using Glmnet(Friedman et al 2010) through Lasso analysis. In this step, isoforms were used as the predictor variable (x) and fiber development stage as the response variable (y), with 5-fold cross-validation. The variables selected at Lambda.min were prioritized for subsequent analysis, leading to the identification of key isoforms associated with fiber development.

### Species-Specific Isoform Identification

First, the longest transcript’s protein sequence was extracted to represent each gene, followed by orthologous gene identification between 3-79 and TM-1 using OrthoFinder(Emms and Kelly 2019). Transcript and CDS sequences were the prepared, along with the corresponding annotation files. EGIO (Exon Group Ideogram-based detection of Orthologous Exons and Orthologous Isoforms)(Ma et al 2022) was used to identify orthologous isoforms between the two species.

## Conclusion

This study constructed the most complete and comprehensive transcriptome atlas of *Gossypium barbadense* and *Gossypium hirsutum*, identifying alternative splicing events across 10 fiber developmental stages. It also identified isoforms most relevant to fiber development, providing a theoretical foundation for improving cotton fiber quality traits.

## Discussion

Based Iso-seq to characterize full-length isoform construct the comprehensive isoform-resolved cotton transcriptome by combing short read RNA-seq and generate elaborate landscape of AS in the *Gossypium barbadense* and *Gossypium hirsutum*. Surprisingly, it is discovered that there is significant isoform diversity among genes in both 3-79 and TM-1, including a considerable number of novel isoforms do not present in existing genome annotations, particularly in genes expressed during the fiber developmental stages.

The current gene annotations are incomplete, and a considerable proportion of genes contain numerous novel isoforms. By utilizing Iso-seq to refine the existing gene annotations, previous Transcription Start Sites and Transcription Termination Sites have been corrected. Iso-seq-derived TSSs have proven to be more accurate than the existing TSSs, as confirmed by the patterns of accessible chromatin observed in ATAC-seq data. Further in-depth exploration is still required in the future.

Genes associated with fiber developmental stages are characterized by considerable isoform diversity, suggesting that these novel isoforms play a vital role in growth.

## ACKNOWLEDGMENTS

The computations in this paper were run on the bioinformatics computing platform of the National Key Laboratory of Crop Genetic Improvement, Huazhong Agricultural University.

## Data availability

The datasets used in this study are publicly available from the following repositories:

PRJNA726938

PRJNA359724

PRJNA503814

CNP0002656

PRJNA576032(ATAC-seq)

PRJNA396502(DNase-seq)

## Code availability

The Quarto files, along with related scripts, have been uploaded to GitHub (https://github.com/Tang-pro).

## Notes

### Competing Interest Statement

The authors have declared no competing interest.

